## Supplementary figures and images for "Host population structure and rare dispersal events drive leptospirosis transmission patterns among *Rattus norvegicus* in Boston, Massachusetts, US"

### Supplemental Figure 1

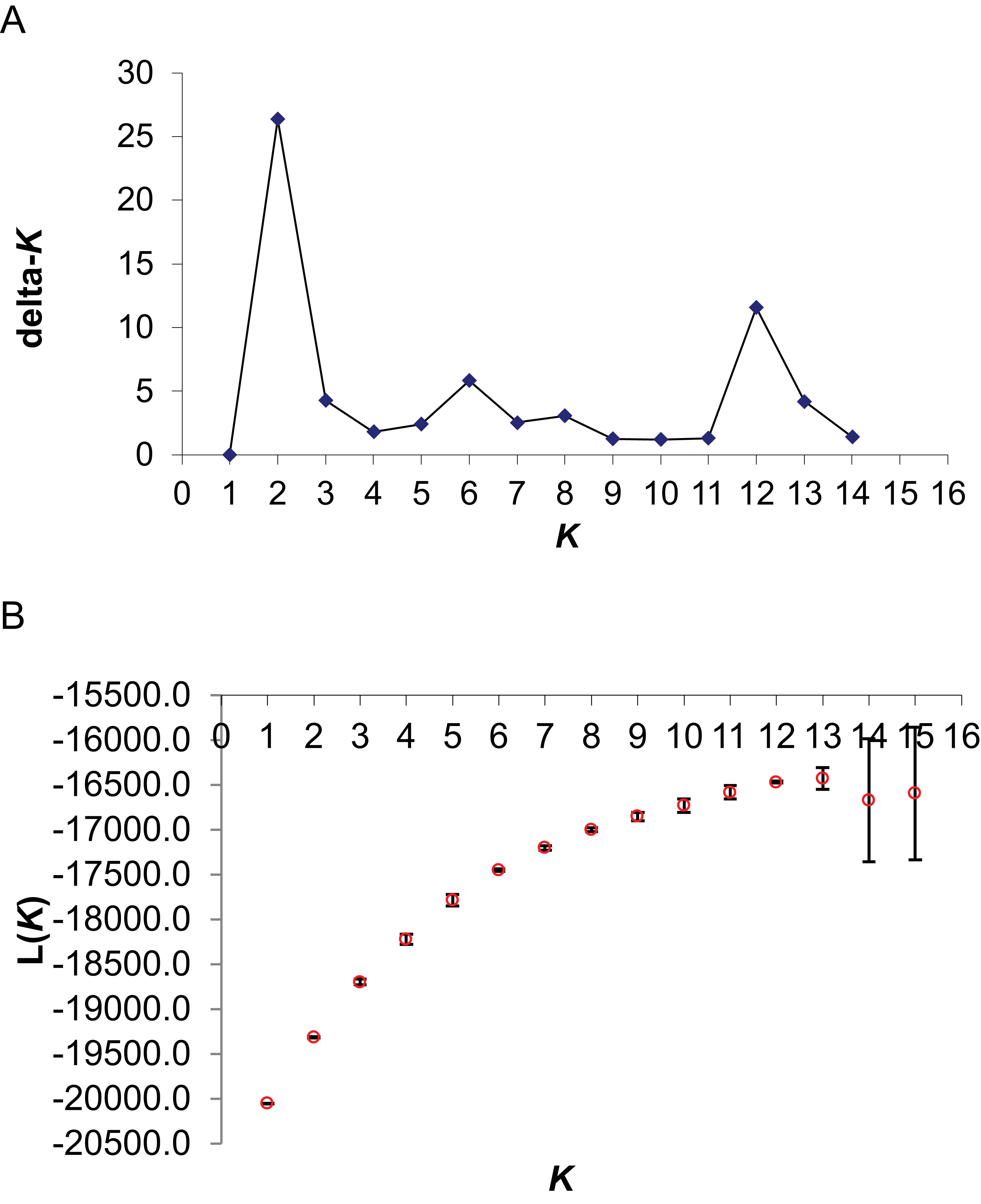

### Supplemental Figure 2

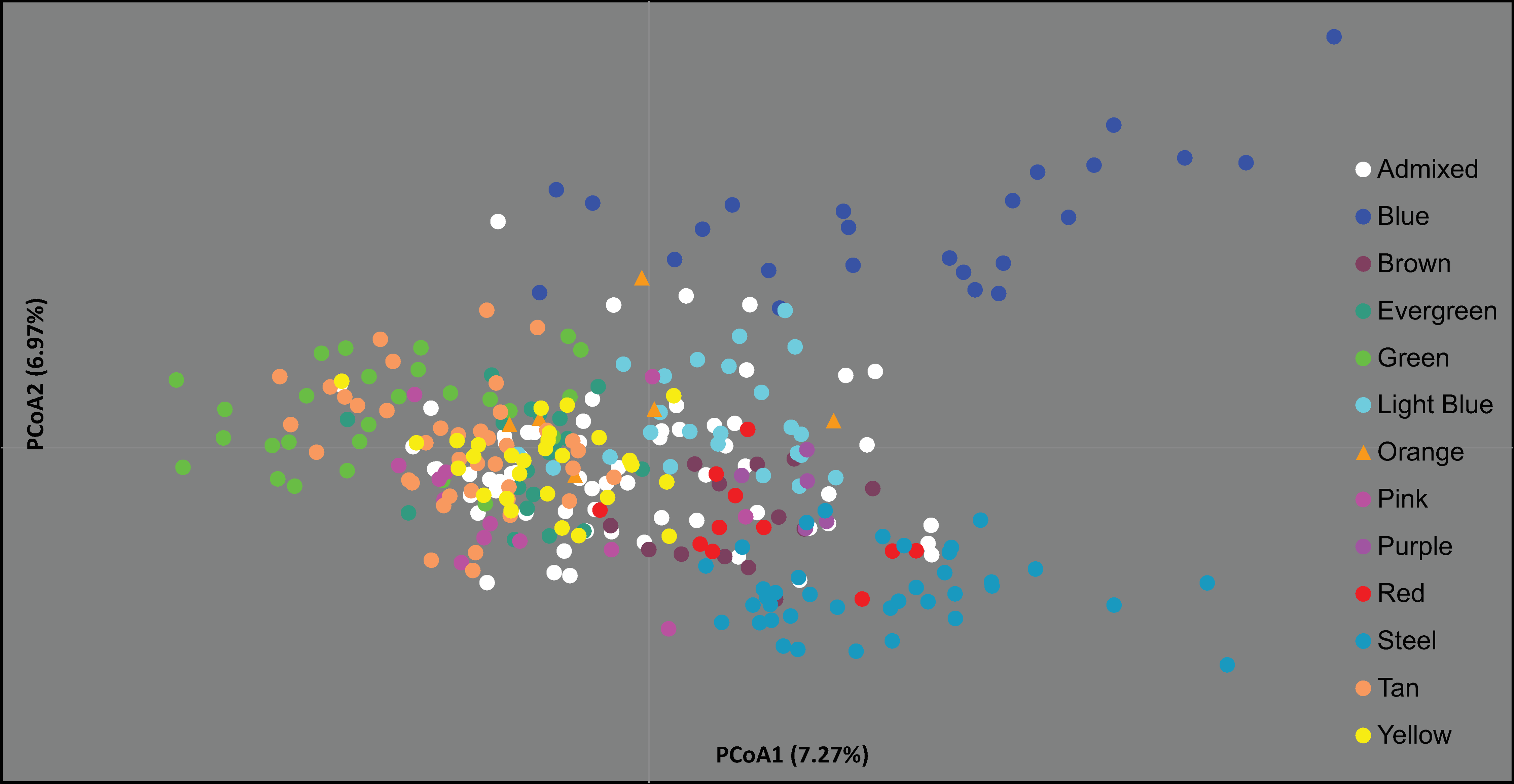

### Supplemental Figure 3

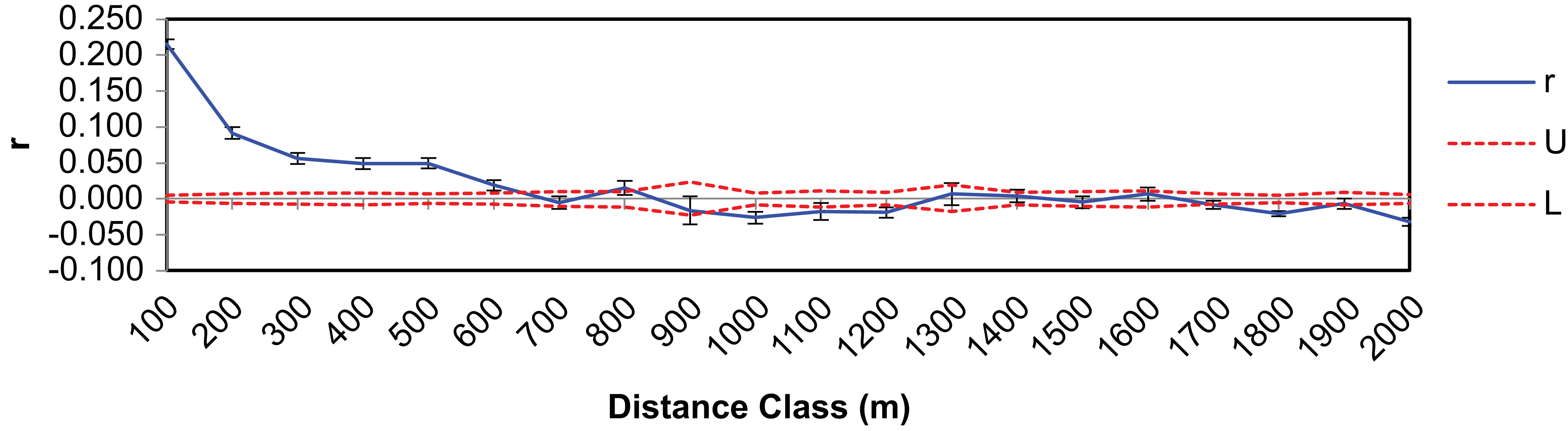

### Supplemental Figure 4

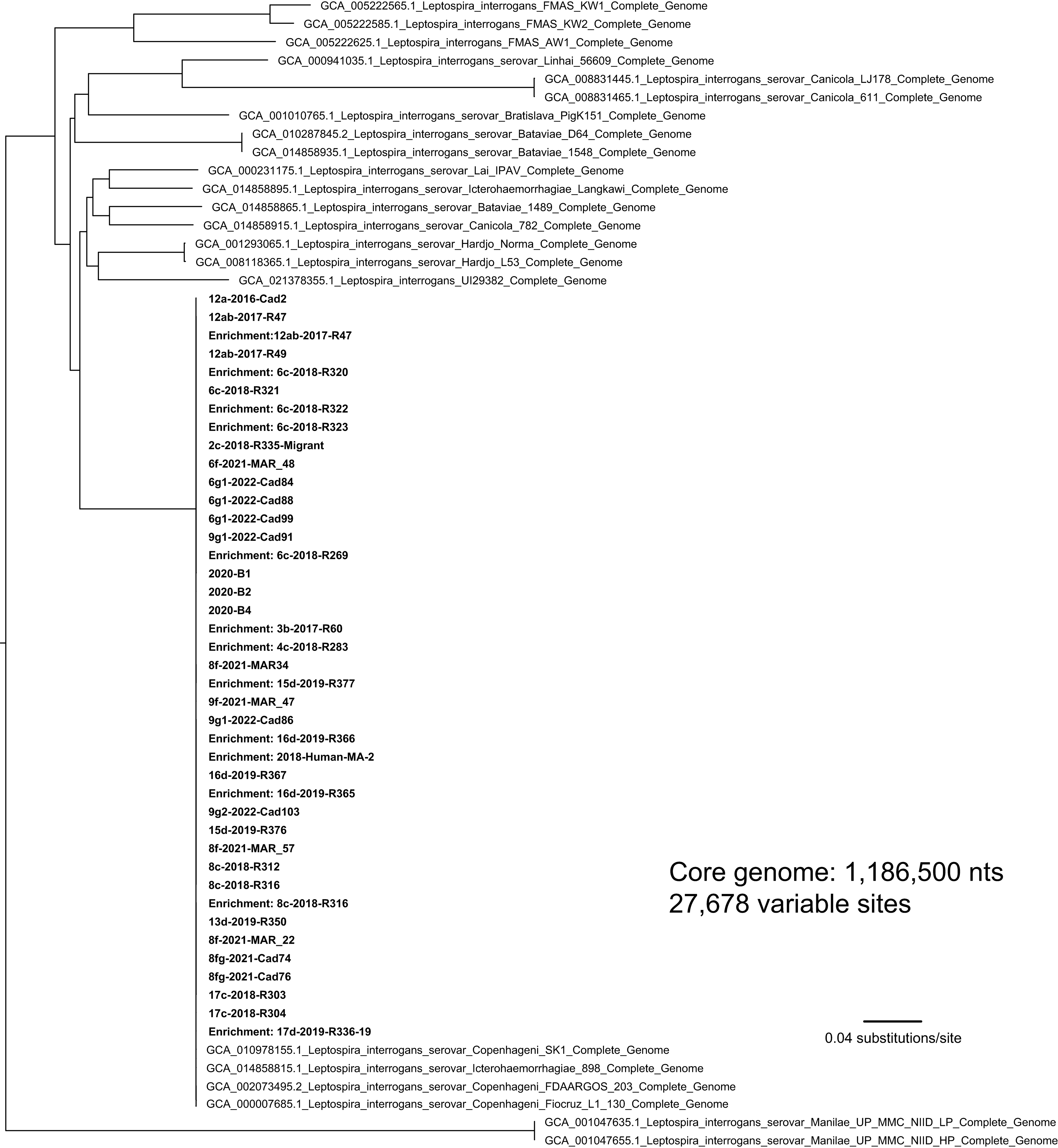

### Supplemental Figure 5

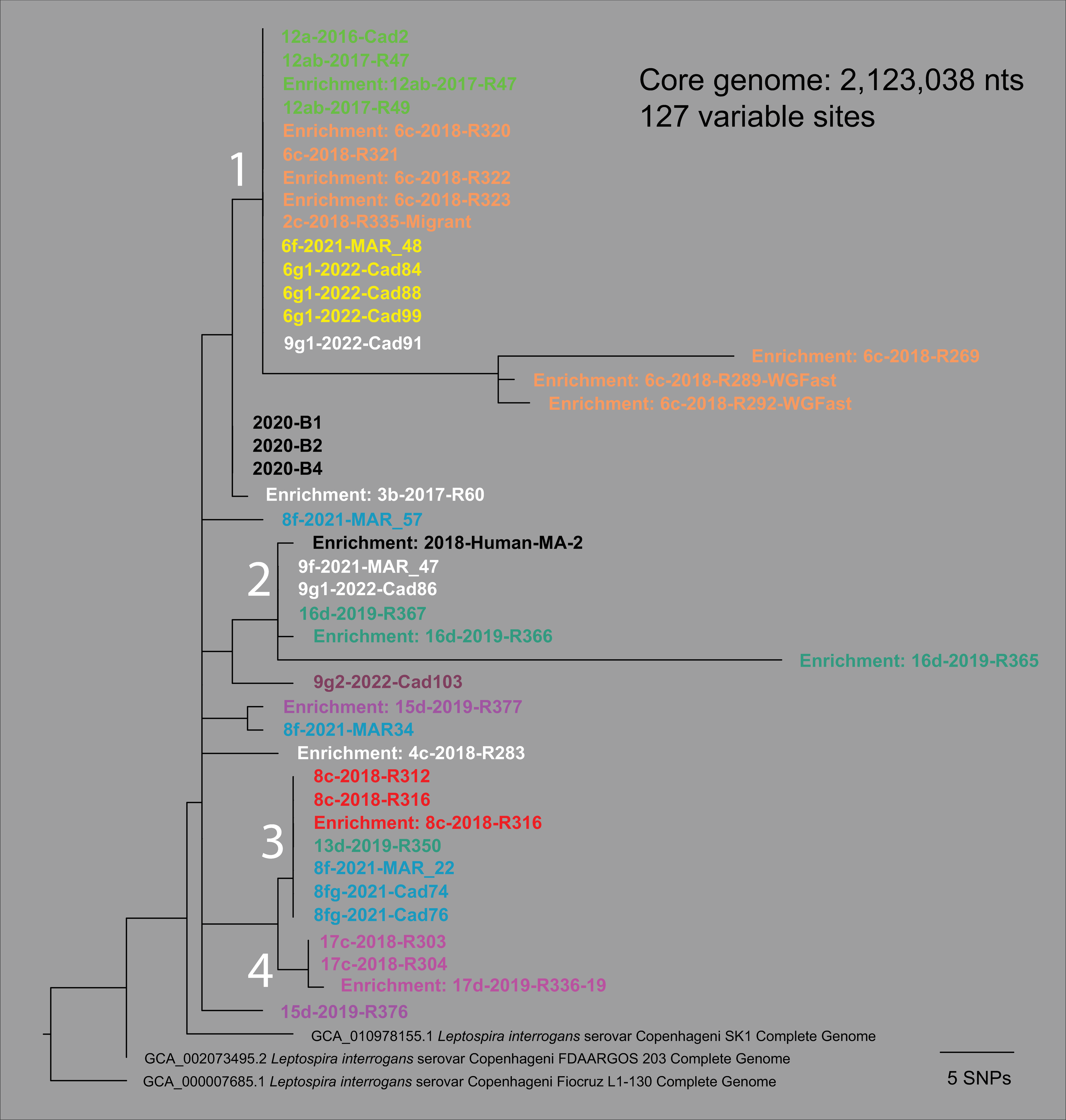

### Supplemental Figure 6

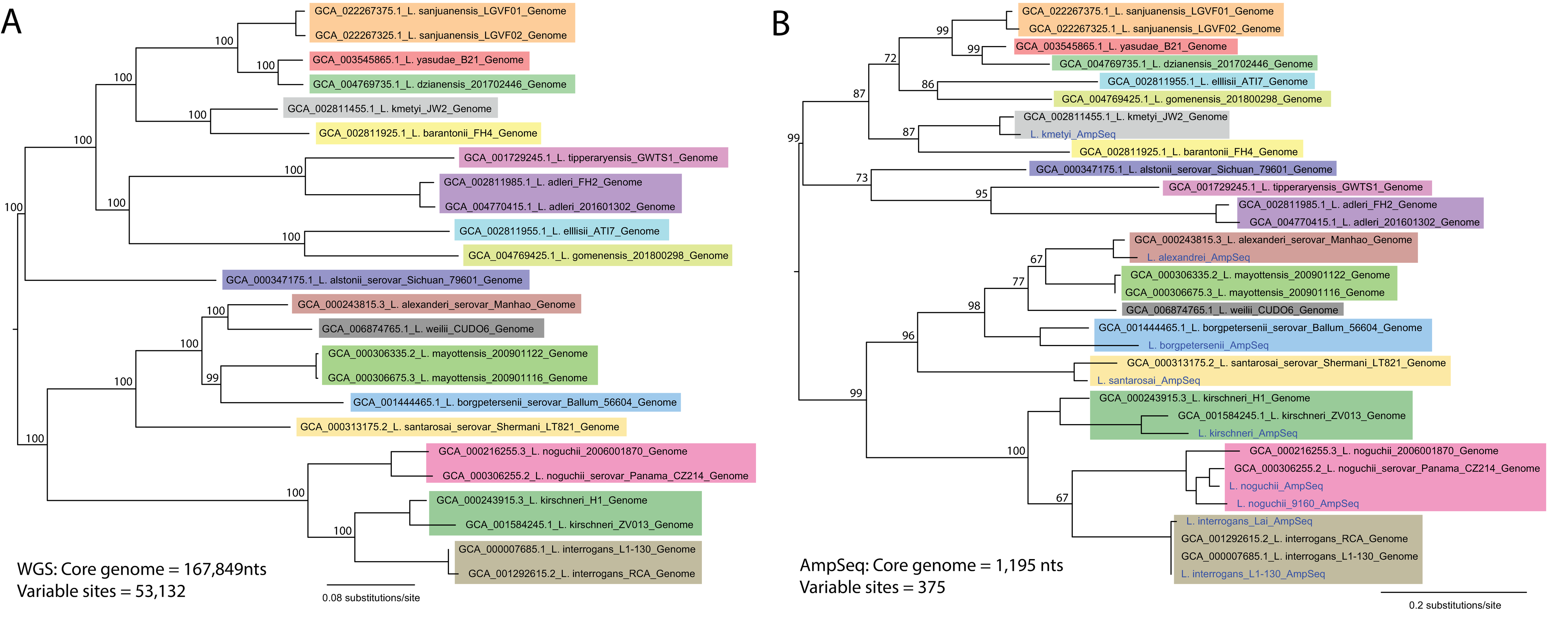

### Supplemental Figure 7

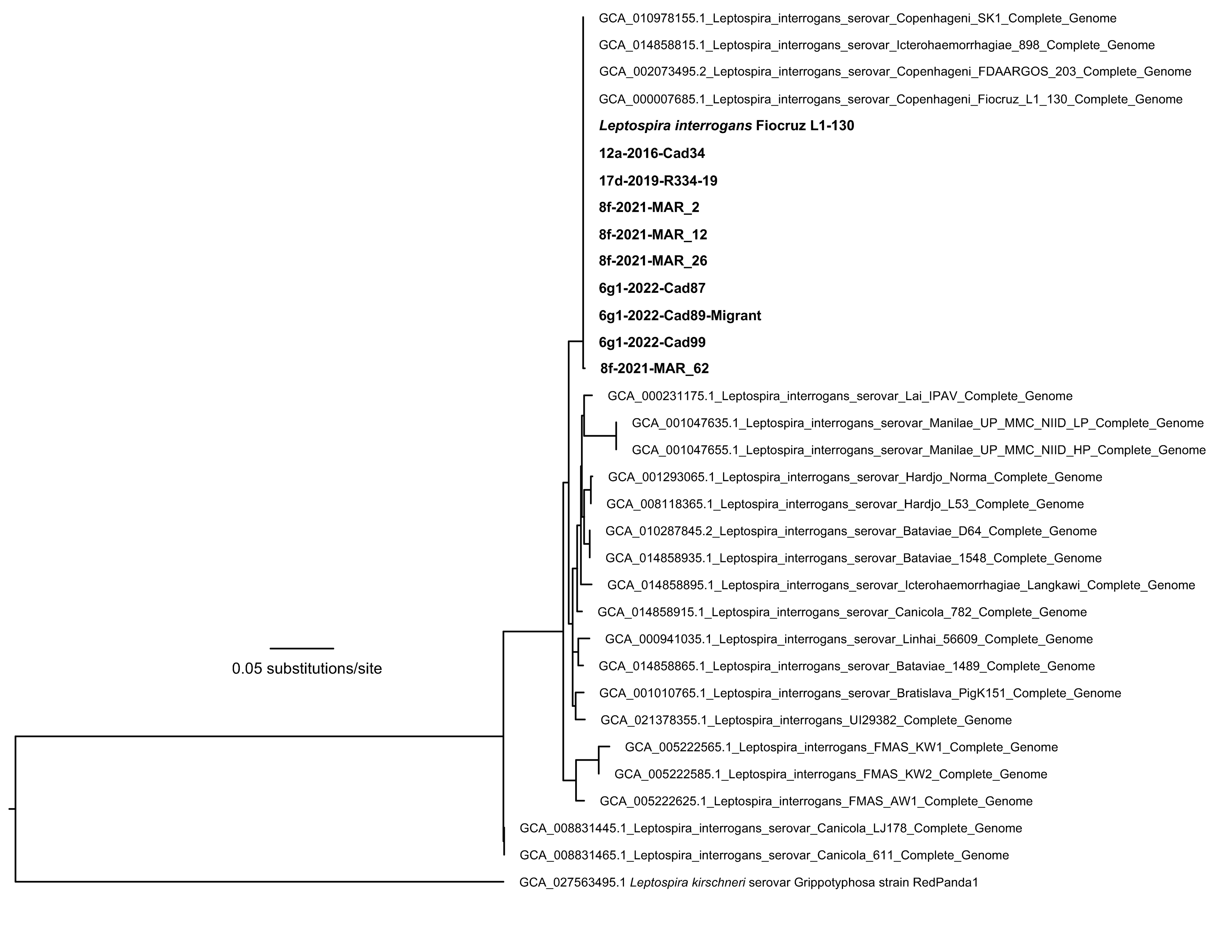
